# Activity-dependent decline and recovery of synaptic transmission in central parts of surviving primary afferents after their peripheral cut

**DOI:** 10.1101/2022.06.29.498096

**Authors:** Morgane Le Bon-Jégo, Marie-Jeanne Cabirol, Daniel Cattaert

## Abstract

Whereas axons deprived of their nucleus degenerate within a few days in Mammals, they survive for several months in Crustacean. However, it is not known if central synapses from sensory axons may preserve their molecular machinery in the absence of spiking activity, after peripheral axotomy, which suppress their nucleus. Using electrophysiology techniques and electron microscopy imaging we report that 1) Electron microscopy analysis confirms previous observations that glial cell nucleus present in sensory nerve, proliferate an migrate to axon tubes, in which they form close contact with surviving axons; 2) after peripheral axotomy performed *in vivo* on Coxo-Basipodite chordotonal organ (CBCO) sensory nerve does not convey any sensory message, but antidromic volleys are observed; 3) Central synaptic transmission to motoneurons (MNs) progressively declines over #200 days (90% of monosynaptic excitatory transmission is lost after 3 weeks, whereas 60% of polysynaptic inhibitory transmission persist up to 6 months). After #200 days no transmission is observed anymore; 4) However, this total loss is only apparent, because repetitive electrical stimulation of the sensory nerve *in vitro* progressively restores first inhibitory post-synaptic potentials (IPSPs) then excitatory post-synaptic potentials (EPSPs); 5) The loss of synaptic transmission can be prevented by *in vivo* chronic sensory nerve stimulation; 6) Using simulations based on the geometric arrangements of synapses of the monosynaptic excitatory transmission and disynaptic inhibitory pathways, we have shown that antidromic activity in CBCO nerve could play a role in maintenance of synaptic function of inhibitory pathways to MNs, but not on monosynaptic excitatory transmission to MNs. Taken together, our study confirms the key role of glial nucleus in axon survival, that machinery for spike conduction and synaptic release even if no activity is present for several months. After long silence periods (>6 months) spike conduction and synaptic function can still be restored by electrical activity.

## Introduction

Contrary to Vertebrates in which axons do not survive when deprived of their nucleus (Koeppen, 2004), the peripheral part of cut soma-deprived motor axons do not degenerate in crustaceans thanks to invading glial soma (Bittner & Baxter, 1991; Parnas *et al*., 1998). This distal axon survival allows for their functional reconnection to the outgrowing proximal stump of the motor axon (Hoy *et al*., 1967), as indicated by regenerative sprouting response observed both in motor peripheral and sensory central cut ends (Kennedy & Bittner, 1974). Moreover, during the first months, the soma-deprived motor axons keep their capacity to conduct spike and trigger transmitter release (Bittner, 1973). The same observations were made for sensory axons which have their cell bodies located in peripheral sensory organs. Once cut, the soma-deprived central axons maintain their spike conduction and synaptic release capacities, and this maintaining is accompanied by glial cell reorganization too (Govind *et al*., 1992).

Peripheral motor axons of cut motor nerves continue to conduct spikes for at least a year after they are cut (Atwood *et al*., 1989), with a velocity that is not different from control axons (Parnas *et al*., 1991). However, even though such cut axons continue to elicit neurotransmitter release even one year after cut, synaptic release kinetics undergoes deep changes along time. In particular, the decay phase of quantum events that followed a single exponential decay in control preparations, are much slower in cut axons and can no more be fitted with one exponential decay. Moreover, the duration of the synaptic event increases progressively after the section. The longer decay phase of postsynaptic events is probably explained by post-synaptic mechanisms (Parnas *et al*., 1991) while the prolongation of the time course is likely presynaptic.

Concerning sensory axons, their survival and degeneration after nerve cut, was studied in crayfish tailfan sensory nerves (Govind *et al*., 1992). Contrary to motor axons, in this model, 95% of sensory axons deprived of their cell body, underwent a degenerative process (Govind *et al*., 1992). Central connections of remaining axons onto target neurons, was deeply modified. As was observed in motor axons, surviving sensory axons were invaded by glial cell nuclei (Govind *et al*., 1992).

In axons that have been cut from their cell body (i.e peripheral stumps for motor axons, and central parts for sensory axons), the lack of spike for long periods of time (up to one year) could potentially lead to dramatic changes in synapse function. Indeed, activity-dependent changes in synaptic function has been shown in invertebrates (Lnenicka, 2020; Goel & Dickman, 2021) and vertebrates (Magee & Grienberger, 2020). In Hebbian rules of synaptic plasticity, when a presynaptic neuron is not involved in the activity of the post-synaptic neuron, its synaptic transmission decreases. If such mechanisms are present, lack of activity could be involved in the synaptic changes observed in motor (Parnas *et al*., 1991) and sensory (Govind *et al*., 1992) axons deprived of their cell body and source of activity.

In this paper, we addressed this question by studying the synaptic transmission in a well-documented sensorimotor circuit of the crayfish composed of the coxa-basis chordotonal organ (CBCO) which contains sensory neurons ensuring the resistance reflex via monosynaptic connections with depressor and levator MNs (El Manira, Cattaert *et al*., 1991; El Manira, DiCaprio *et al*., 1991), and disynaptic inhibitory connections with antagonistic MNs (Morgane Le Bon-Jego & Cattaert, 2002). After *in vivo* section of the sensory CBCO nerve, the sensorimotor system was dissected out after various periods of time (from a few days to more than 6 months post lesion) to be studied *in vitro*. This allowed to follow the evolution of monosynaptic excitatory synaptic transmission and disynaptic inhibitory transmission. In parallel, electronic microscopy studies were achieved on the central part of the cut sensory nerve to confirm the maintenance of CBCO axons over months, and the involvement of glial cells. In a series of experiments, we show that all synaptic transmission is lost 3 months after peripheral cut of the CBCO nerve. In order to test if this loss was due to lack of sensory activity, we repeated these experiments, in which we placed two cuff stimulating electrodes on the central end of the cut nerve, allowing to maintain orthodromic spiking activity during the period of time following the cut of the sensory nerve. We also show that, even in the absence of sensory activity for long periods of time *in vivo*, it is still possible to re-establish functional synaptic connections with MNs and with inhibitory interneurons of the disynaptic pathway.

## Material and methods

### Experimental animals

Experiments were performed on adult crayfish (*Procambarus clarkia*) of either sex measuring about 12 cm. The animals were purchased from a commercial supplier (Chateau Garreau, France), and maintained in indoor aquaria at 15-18°C. Animals were fed once a week. After surgery, the animals were kept in individual aquaria.

### In vivo surgery, stimulation and recordings

The sensory nerve of the primary afferent is very superficial allowing its section *in vivo* on anesthetized animals (as previously described in (Le Ray *et al*., 2005)) and the implantation of extracellular electrodes. The procedure used for chronic recordings was adapted from the technique developed by (Böhm H., 1996). Animals were anesthetized on ice, and immobilized (as previously described in (Le Ray *et al*., 2005) for the dissection.

#### In vivo section of the CBCO sensory nerve

A small piece of cuticle from the coxo-basipodite joint of the 4th walking leg was removed to access to the CBCO sensory nerve and to section it. After the efficiency of the section was confirmed by the disappearance of the resistance reflex (Fig. 1A1-2), the hole in the cuticle was filled with a mixture of colophane and heated bee wax. Animals were then placed in individual aquaria until used for recordings or electron microscopy.

**Figure 1.**
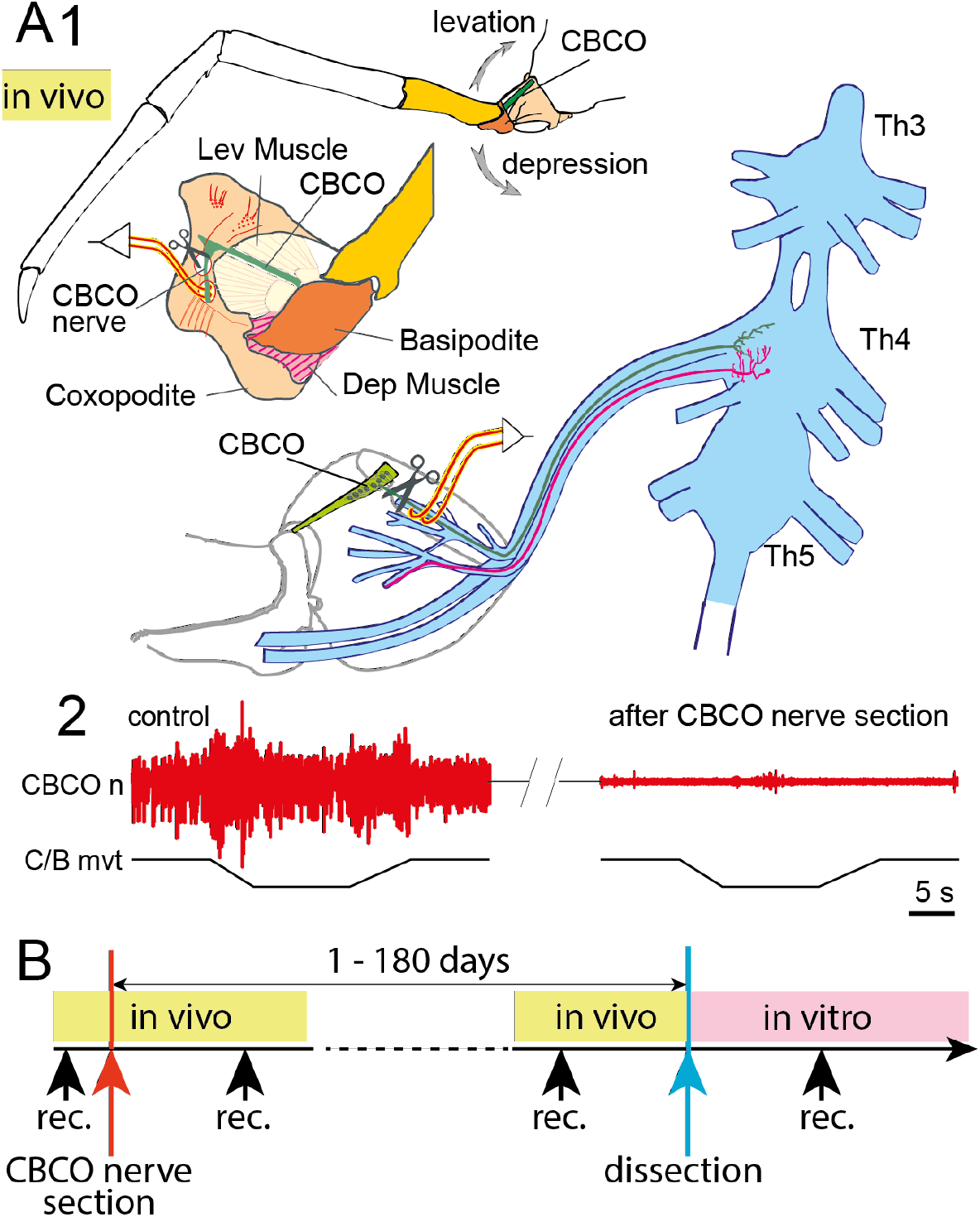
(methods): Description of the sensory-motor system. **A1:** Organization of the coxo-basipodite joint of the crayfish walking leg controlling upward and downward movement of the leg. A proprioceptor, the coxo-basipodite chordotonal organ (*CBCO* in green), encodes the vertical movements of the leg. An *in vitro* preparation of the locomotor nervous system (not shown) can be obtained, it consists of thoracic ganglia (*Th3-Th5*) dissected out together with motor nerves of the proximal muscles (*Lev* and *Dep*) and the sensory nerve. This preparation allows extracellular and intracellular recordings of the different element of the network. The CBCO sensory nerve was cut *in vivo*. Cuff recording electrodes were used to record activity from the CBCO nerve (see methods) on the central part of the nerve before (A2, left) and after (A2, right) section of the nerve. **A2**: Electroneurogram of the sensory nerve (*CBCO. n*) showing the effective section of the sensory nerve. **B**: Timeline of the experiments. In the various experiments, CBCO nerve could be recorded before section, and from 1 to 180 days after section *in vivo* (see yellow part of time-line). Dissection of the operated crayfish was performed after various delays after CBCO nerve section *in vivo* to perform intracellular recordings *in vitro* (see pink part of the timeline).

#### In vivo recordings and chronic stimulation of the sensory nerve

Four thin monopolar wires, traveling in a grounded cable fixed with wax on the back of the animals were used. Three of them were attached to 50 µm insulated wire electrodes to record muscular and nerve activities (Fig. 1), whereas a 4^th^ electrode was placed under the carapace and used as a reference. Electrodes were isolated from the hemolymph with a flexible silicone elastomer (World Precision Instruments). Each recording electrode was fixed separately on the leg cuticle with wax before it reached its attachment to the grounded cable and connection to homemade extracellular amplifiers. Amplified neurograms and EMG signals were directed through a CED 1401 interface (Cambridge Electronic Design) to a computer for storage and analysis. For the chronic stimulation of the sensory nerve, trains of 500 ms duration were applied at a frequency of 0.25 Hz. Each train was composed of 0.3 ms pulses at 20 Hz.

### In vitro preparation and recordings

#### In vitro preparation

After various delays (from 1 day to 180 days after CBCO nerve section) (see protocol in Fig. 1B), the ventral thoracic nerve cord was dissected out with the sensory and motor nerves to the 4^th^ left leg. This *in vitro* preparation was used to study sensory-motor connections ((El Manira, Cattaert *et al*., 1991; Morgane Le Bon-Jego & Cattaert, 2002; M. Le Bon-Jego, 2004; Morgane Le Bon-Jego *et al*., 2006). It consists of the last three thoracic (*T3-T5)* ganglia dissected out along with all the nerves (motor and sensory) of the two proximal segments of the left fourth leg (Fig. 1). In control experiments, the chordotonal organ (CBCO), which monitors the movements of the second joint (coxo-basipodite), was also dissected out and kept intact. For the operated, the remaining part of the sensory nerve without the CBCO was carefully dissected. The preparation was pinned down dorsal side up in a silicone elastomer covered petri dish. The fourth ganglia were desheathed to improve the continuous superfusion of the central neurons with oxygenated saline and to allow intracellular recordings of all Dep MNs. The saline was composed of (in mM) 195 NaCl, 5 KCl, 13 CaCl2, 2 MgCl2, buffered with 3 mM N-2-hydroxyethylpiperazine-N’-ethanesulfonic acid (HEPES, Sigma Chemical) and pH adjusted to 7.65 at 15°C. In some experiments, in order to rise the spiking threshold of interneurons, we used a high divalent cation solution containing (in mM) 34 CaCl2, 6.4 MgCl2 with sodium concentration reduced accordingly to preserve the osmolarity of the solution (157 NaCl). The use of this altered saline allowed us to identify the monosynaptic reflex responses (Berry & Pentreath, 1976).

#### Ex vivo recordings

**Extracellular recordings** from the motor nerves innervating the depressor and levator muscles and from the sensory nerve of the CBCO were performed using stainless steel pin electrodes contacting the nerves and insulated from the bath with Vaseline. Recorded signals were amplified by differential AC amplifiers (Grass, Quincy, MA, gain of 10 000×). **Intracellular recordings** from depressor MNs (Fig. 4) were performed from their main neurite within the ganglion using glass micropipettes (Clark Electromedical Instruments, Reading, UK) filled with 3 M KCl (resistance 20-25 MΩ) connected to an Axoclamp 2B amplifier (Axon Instruments Inc, Foster City, CA) used in the current-clamp mode. Dep MNs were identified using the following criteria: 1) the spikes evoked by the electrical stimulation of the Dep nerve was recorded by the intracellular microelectrode and 2) the spikes evoked by a depolarizing current injected into the intracellularly recorded neuron were correlated one-to-one with the extracellular spikes recorded in the Dep nerve. An eight-channel stimulator (A.M.P.I., Jerusalem, Israel) was used for intracellular stimulation of MNs during the identification procedure, and for CBCO nerve stimulation. Data were digitized and stored onto a computer hard disk through an appropriate interface (1401plus) and software (Spike2) from Cambridge Electronic Design Ltd. (Cambridge, UK).

### Study of the functional connections

The changes of the sensory-motor connections after section of the CBCO sensory nerve were tested *in vitro*. The CBCO nerve (*nCBCO* s*tim* in Fig. 4) was electrically stimulated and the evoked response in the Dep MN was recorded intracellularly (Fig. 4A). Classically, this electrical stimulation protocol activates simultaneously the two components of the resistance reflex (monosynaptic excitation – *EPSP* - and disynaptic inhibition – *IPSP*) because release and stretch sensitive CBCO neurons are located in the same nerve (see schema in Fig.4A1).

### Data analysis

Data were analyzed using the Spike2 analysis software. Statistical analyses were performed with Prism (Graphpad Software, San Diego, CA). Results are given as mean values ± standard error of the mean (SEM). The significance of the effect of peripheral axotomy on the synaptic connection (number of MN eliciting an EPSP or an IPSP) along time was assessed by One Way ANOVA followed by Tukey’s multiple comparison tests (see Fig. 4B4, right). Comparison between evolution of EPSPs and IPSPs along time was assessed by Two Ways ANOVA (see Fig. 4B4, left). Changes in synaptic delays after a given period after axotomy was assessed by unpaired t-tests.

### Electron microscopy

The CBCO sensory nerve from control and operated animals was collected and fixed overnight at 4°C in glutaraldehyde (0.1%). The samples were then rinsed in phosphate buffer containing sucrose, post-fixed at room temperature in osmium 1% for 45 min and stained with uranyl acetate (2%). Dehydration was obtained through an ascending series of ethanol bathes. Nerves were then embedded in araldite and resin polymerized at 60°C for 3 days. Ultrathin sections (70-80 nm) were collected on nickel grids contrasted with uranyl acetate and lead citrate. They were then observed with a transmission electronic microscope (Hitachi H600). For some experiments, grids were immunostained for GABA following the protocol in Watson et al. (2005). Briefly, sections were treated at room temperature with 2% periodic acid (3 minutes) and in sodium metaperiodate at 50°C (3 minutes), then washed in Tris buffer (TB), pH 7.2. After 30 minutes in 5% normal goat serum diluted in TB, the grids were transferred for 2 hours in polyclonal rabbit anti-GABA at 1:1,000 (Sigma) in TB. After further washing, they were transferred for 1 hour to 15 nm gold-labelled goat anti-rabbit antibody (British Biocell International, Cardiff, United Kingdom) diluted 1:20 in TB, pH 8.2, for 1 hour. Finally, sections were counterstained with uranyle acetate and lead citrate. Preabsorption with GABA conjugated to bovine serum albumin (BSA) with glutaraldehyde eliminated all labelling with GABA antibody.

#### Image analysis

The surface occupied by glial cell was calculated as follows. Using ImageJ the contours of glial cells were manually drawn on EM image. The area and perimeter of the contours were then calculated by ImageJ, using the scale bar, and the total image area was used to calculate the percentage of surface occupied by glial cells.

### Computer modeling

In this study, we have modeled the interaction between a GABA synapse responsible for PADs (primary afferent depolarization responsible for presynaptic inhibition) onto disynaptic inhibitory circuit of reciprocal inhibition (Morgane Le Bon-Jego & Cattaert, 2002; M. Le Bon-Jego, 2004). The GABA synapse producing PADs is located on a 1^st^ order branch in the region of the first branching point of a primary afferent (CBCO) sensory fiber in ganglion neuropile (D. Cattaert *et al*., 1992; Daniel Cattaert & El Manira, 1999; Daniel Cattaert *et al*., 2001). The synapses from CBCO to the inhibitory interneuron of the reciprocal inhibition are located on CBCO main axon (Fig. 2A-B). The effect of PADs was simulated using NEURON 7.3 program (Hines and Carnevale 1997). The temporal integration step was 20 µs.

**Figure 2.**
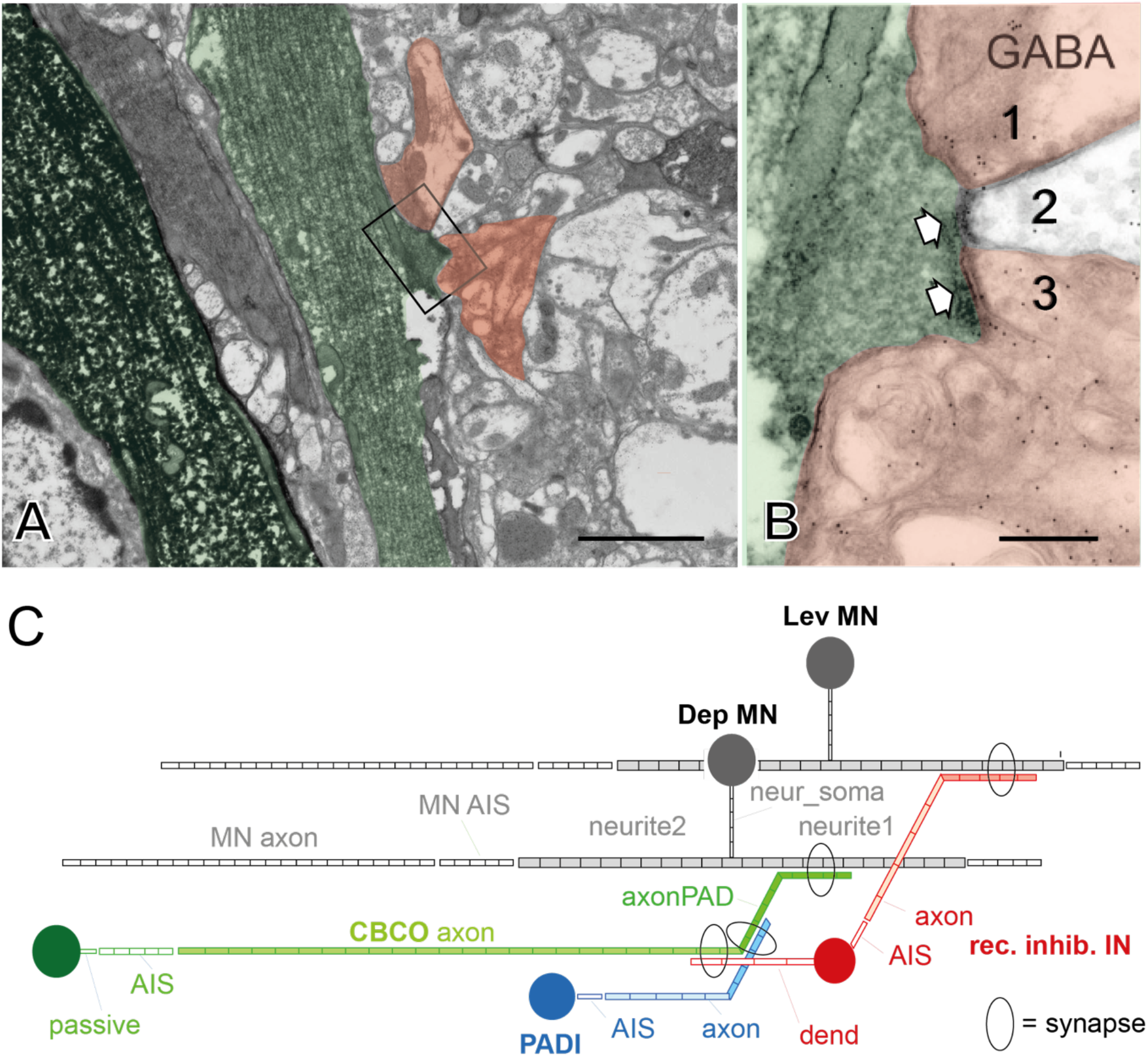
(methods): Description of the five neurons modeled in NEURON script: **A:** Electron microscopy images of two CBCO afferent terminals, one lightly stained and the other heavily stained. The darkly stained one runs within the nerve root, where it is isolated from the neuropil, but the other one has emerged from the tract. The region indicated by the box is seen at higher magnification in B. **B**: A small swelling on the axon packed with agranular vesicles makes two synapses (white arrows) onto three postynaptic processes (1–3), which contain a small number of agranular vesicles. Two of these processes (1 and 3) appear to be immunoreactive for GABA. A postsynaptic electron-dense thickening of the membrane is seen in each of the processes. **C: Compartment models used in Neuron simulation. Four neurons were modeled** ***1) Motoneuron (MN* in grey)**. The *<u>soma</u>*, which does not participate to electrical activity of the MN is connected to the main neurite via a thin neurite (*neur_soma*). The main neurite is made of two sections: *<u>neurite 1</u>*, in which is located the post-synaptic compartment of the synapse from the reciprocal inhibition neuron (*rec. inhib. IN* in red) *and <u>neurite 2</u>*, in which is located the post-synaptic compartment of the synapse from the CBCO afferent neuron *(CBCO axon* in green) ***2) CBCO afferent neuron (CBCO* in green)**. Composed of seven sections: *<u>soma</u> i*n which current is injected to produce a spike; *<u>passive section (passive)</u>* connecting the soma to the axon initiating spike section (AIS); *<u>axon initial segment (AIS)</u>* in which sensory spikes are normally triggered (but not in this simulation); *<u>axon (CBCO axon)</u>* conveying actively spikes to CNS; *<u>active section</u>* with output synapse to the reciprocal inhibitory interneuron *(rec. inhib. IN); <u>passive section</u> r*eceiving inhibitory synapse from the primary afferent depolarization interneuron *(PADI* in blue) producing PADs in the primary afferent terminal, and antidromic spikes conveyed in the axon *to the periphery; <u>passive section</u>* conveying depolarization to the excitatory synapse to the MN neurite; *<u>passive section</u>* where the output excitatory synapse to the MN is located. ***3) Primary Afferent Depolarization Interneuron (PADI* in blue)** *c*omposed of a *<u>soma</u>*, an *<u>axon initial segment (AIS)</u>*, an *<u>axon</u>* and an *<u>active section</u>* where output inhibitory synapse to the CBCO neuron is located. Spikes are triggered by current injection in the soma (see Fig. 5D). ***4) Reciprocal inhibitory Interneuron (rec. inhib. IN* in red)** composed of four sections: *<u>dendrite</u>* receiving synaptic input from the CBCO Neuron; *<u>soma</u>*; *<u>axon initial segment (AIS)</u>* in which sensory spikes are produced; *<u>axon</u>* conveying spikes actively; *<u>active section</u>* with output synapse onto MN neurite 1 See table for the conductance equipment of each section. Figures 2A and 2B are modified from (Watson *et al*., 2005).

Five neurons were simulated: a primary afferent fiber and terminal, (in green in Fig. 2C and Fig. 8A) a MN, a PADI (inhibitory neuron producing PADs in the CBCO terminal, in blue in Fig. 2C and Fig. 8A), and a reciprocal inhibition interneuron (in red in Fig 2C and Fig. 8A). Each neuron was wade of several parts (soma, axon initial segment, axon, and synaptic compartments (either for input or output). The two antagonistic MNs (Dep and Lev) were more complex, with some passive sections (soma, thin neurite between soma and large neurite – neurite1 and neurite2) and some active sections (AIS and axon). Note that input synapses on MN were placed onto neurite1 (see Fig 2C). The primary afferent conveyed spikes in axon but not in the terminal branches. The dimensions and properties of all neuron compartments are given in table 1.

**Table 1.** Primary afferent neuron

|  | soma | passive | AIS | axon | PAD_area | terminal |
| --- | --- | --- | --- | --- | --- | --- |
| diam (um) | 10 | 2 | 2 | 1 | 1 | 1 |
| length (um) | 10 | 10 | 30 | 500 | 50 | 50 |
| Rm (Ohm.cm2) | 3000 | 3000 | 3000 | 3000 | 3000 | 3000 |
| Ra (Ohm.cm) | 100 | 100 | 100 | 100 | 100 | 100 |
| gmax_Na (uS) | 0 | 0 | 0.8 | 0.09 | 0.03 | 0.03 |
| gmax_K (uS) | 0.005 | 0.005 | 0.075 | 0.036 | 0.016 | 0.016 |

**Motoneuron**
|  | soma | dend_soma_MN | neurite1_MN | MN_syn_in_area | neurite2_MN | AIS_MN | axon |
| --- | --- | --- | --- | --- | --- | --- | --- |
| diam (um) | 40 | 1 | 5 | 2 | 5 | 2 | 2 |
| length (um) | 40 | 100 | 100 | 20 | 100 | 30 | 500 |
| Rm ($\Omega$.cm2) | 3000 | 3000 | 3000 | 3000 | 3000 | 3000 | 3000 |
| Ra (Ohm.cm) | 100 | 100 | 100 | 100 | 100 | 100 | 100 |
| gmax_Na (uS) | 0 | 0 | 0 | 0.03 | 0 | 0.8 | 0.24 |
| gmax_K (uS) | 0.005 | 0.005 | 0.0036 | 0.0036 | 0.0036 | 0.075 | 0.036 |

**Reciprocal inhibition interneuron**
|  | soma | dend | AIS | axon | syn_out_area |
| --- | --- | --- | --- | --- | --- |
| diam (um) | 10 | 1 | 2 | 2 | 2 |
| length (um) | 10 | 50 | 30 | 65 | 20 |
| Rm ($\Omega$.cm<sup>2</sup>) | 3000 | 3000 | 3000 | 3000 | 3000 |
| Ra (Ohm.cm) | 100 | 100 | 100 | 100 | 100 |
| gmax_Na (uS) | 0.05 | 0 | 0.5 | 0.12 | 0.12 |
| gmax_K (uS) | 0.036 | 0.036 | 0.075 | 0.036 | 0.036 |

**PAD interneuron**
|  | soma | AIS_PADI | axon_PADI | PADI_syn_out_area |
| --- | --- | --- | --- | --- |
| diam (um) | 10 | 2 | 2 | 2 |
| length (um) | 10 | 30 | 65 | 20 |
| Rm ($\Omega$.cm<sup>2</sup>) | 3000 | 3000 | 3000 | 3000 |
| Ra (Ohm.cm) | 100 | 100 | 50 | 100 |
| gmax_Na (uS) | 0.05 | 0.5 | 0.24 | 0.24 |
| gmax_K (uS) | 0.036 | 0.075 | 0.036 | 0.036 |

Note that Na/K channels were present in CBCO branches but their density did not allow active conduction of spikes.

### Passive properties

All compartments had the same specific membrane resistance (*R*_*m*_ = 3000 Ω.cm^2^). All computations were carried out assuming a specific capacitance, *C*_*m*_ of 1µF/cm^2^ and a specific axoplasmic resistance, *R*_*a*_, of 100 Ω.cm.

### Active Properties

Active compartments were equipped with Na and K Hodgkin Huxley (HH) channels. The equilibrium potential for Na^+^ ions was set E_Na_ = +40 mV. The equilibrium potential for K^+^ ions was E_K_ = -90 mV. The density of Na and K channels in each compartment is given in Table 1.

### Synapses

The activation level of the postsynaptic channel, *s*, ranges from 0 to 1 when the synapse is “closed” or “open”, respectively. The maximum conductance *G*_*syn*_ is fixed, but the value of the actual conductance, *G*_*syn*_.*s*, varies with *s*, that is, when the synapse opens and closes; the kinetics of *s* are controlled by two parameters

#### GABA synapses

tau rise = 1 ms; tau decay = 5 ms.

#### Excitatory synapses

tau rise = 0.2 ms; tau decay = 3.5 ms.

The synaptically induced current that enters the post-synaptic compartment is calculated by

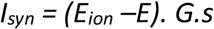

in which *E*_*ion*_ is the equilibrium potential for ions involved in the synapse. For GABA synapses:

#### PADs

*G*_*Cl*_ was fixed to 110 nS and *E*_*Cl*_ was fixed to −35 mV

#### Other GABA synapses

*G*_*Cl*_ was fixed to 0.11 uS and *E*_*Cl*_ was fixed to −70 mV

For Excitatory synapses:

*G*_*Exc*_ was fixed to 22 nS and *E*_*Exc*_ was fixed to 20 mV

## Results

### Proliferation and deep reorganization of glial cell in proximal sensory axons after axotomy

Six months after axotomy, we analyzed the ultrastructure of the sensory nerve. The axons of the central part of the sectioned sensory nerve did not show noticeable change (Fig. 3A,4B). By contrast, glial cells population underwent deep reorganization. Among all profiles in EM, glial cells were identified thanks to their nuclei which are much more electron dense than other profiles. Each nucleus was counted for one cell. Their density increased from 1.37cell/1000µm^2^±0.25 in control to 4.38cell/1000µm^2^ ±1.20 six months after axotomy (Fig. 3H). Their mean individual surface did not change (8.89µm^2^±1.20 in control to 8.62µm^2^±0.60 after axotomy, Fig. 3I). Consequently, the global percentage of surface occupied by glial cell increased from 1.21%±0.28 in control to 3.41%±0.76 after axotomy (Fig. 3J).

**Figure 3:**
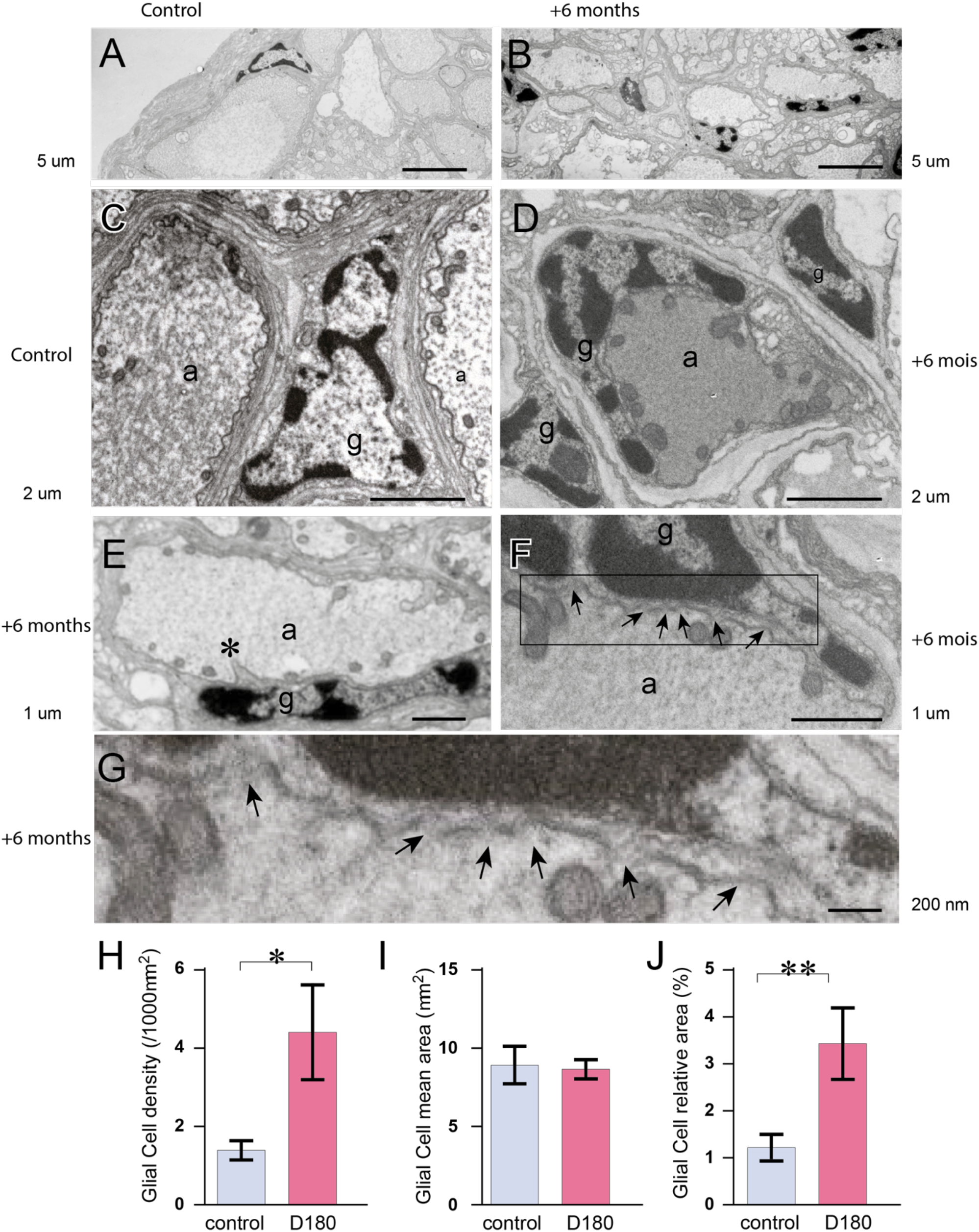
Ultrastructure of the central part of sectioned sensory nerve. **A-B:** Electron microscopy images of control CBCO nerve (*A*) and 6 months (or 180 days, *D180*) cut-CBCO nerve (*B*). Note the increase of glial cell density between control and 180D. **C-D**: Electron microscopy images at higher magnification of control CBCO nerve (*C*) and 6 months (*D180*) cut-CBCO nerve (D). In control (*C*), two neural tubes containing CBCO axons (*a*) and one inter-tube glial cell (*g*) are visible. Six months after CBCO nerve section (*D*), we note two glial cells in inter-tube locations (*g*) and a neural tube containing a CBCO axon (*a*) together with an invading glial cell nucleus (*g*). **E-G**: Electron microscopy images at higher magnification of details from glial cell nucleus and axon relationships in neural tubes presenting some invagination of the glial cell membrane into the CBCO axon (*E*), and discontinuities of the membrane between glial cell and axon in neural tube (see arrows in *F*). The region indicated by the box in *F* is seen at higher magnification in *G*. **H-J**: Statistical analysis of glial cell changes in control (*blue*) and D180 (*pink*). Glial cell density significantly increases in D180 (*H*) while the glial cell mean area is not significantly modified (*I*), but the glial cell relative area significantly increased (J). *: p<0.05; **: p<0.01. Scale bars: 5µm (A, B), 2 µm (C, D), 1 µm (E, F), 200 nm (G).

Glial cell proliferation was accompanied by a change in their anatomical disposition with respect to axons. In control (Fig. 3C) glial cells nuclei were clearly separated from axons, and remained outside of endoneurium. Six months after axotomy glial cells nuclei migrated inside endoneuria, tending to enwrap axons (Fig. 3D). Moreover, the close contact between axons and invading glial cells nuclei, presents anatomical features characterized by invagination (Fig. 3E) and membrane discontinuities (Fig. 3F,G).

### Progressive loss of sensorimotor synaptic transmission from axons deprived from nucleus

After cutting the peripheral sensory nerve from its sensory organ (CBCO) that contains the cell bodies, the sensory axons do not convey any sensory information to the central nervous system, but display sparse antidromic activity (Fig. 7).

To assess the evolution of central synapses from sensory CBCO axons onto target neurons (motoneurons [MNs] and interneurons [INs]) after sectioning the CBCO nerve *in vivo*, dissections were performed after various periods (from a few weeks to > 6 months post CBCO-nerve section, Fig.4B). In these *in vitro* preparations, we electrically stimulated the remaining central part of the sensory nerve. The electrical stimulation of the sensory nerve evoked mixed excitatory/inhibitory responses in intracellularly recorded motoneurons because it stimulated both the direct stretch reflex excitatory pathway (El Manira, Cattaert *et al*., 1991) (Fig. 4A1) and the inhibitory reciprocal innervation circuits (Morgane Le Bon-Jego & Cattaert, 2002). In single experiments, all 12 depressor MNs (Dep MNs) controlling movements of the CB joint (Fig. 4A2) were intracellularly recorded allowing to follow the evolution of identified sensorimotor synapses (Fig. 4B1) along time. In control condition, 9 out of the 12 Dep MNs present monosynaptic EPSP, while 3 of them show only polysynaptic PSPs (Le Ray & Cattaert, 1997) (Fig. 4A). During the three weeks that followed the nerve section, monosynaptic EPSPs progressively declined, and less than 10% of MNs presented an EPSP (Fig. 4B2, 4B3). By contrast, in 60% of MNs, oligosynaptic IPSPs could still be recorded up to 6 months after section (Fig. 4B2, 4B3). After this delay, no PSPs were ever recorded from MNs (N= 3 animals, n=29 MNs).

**Figure 4:**
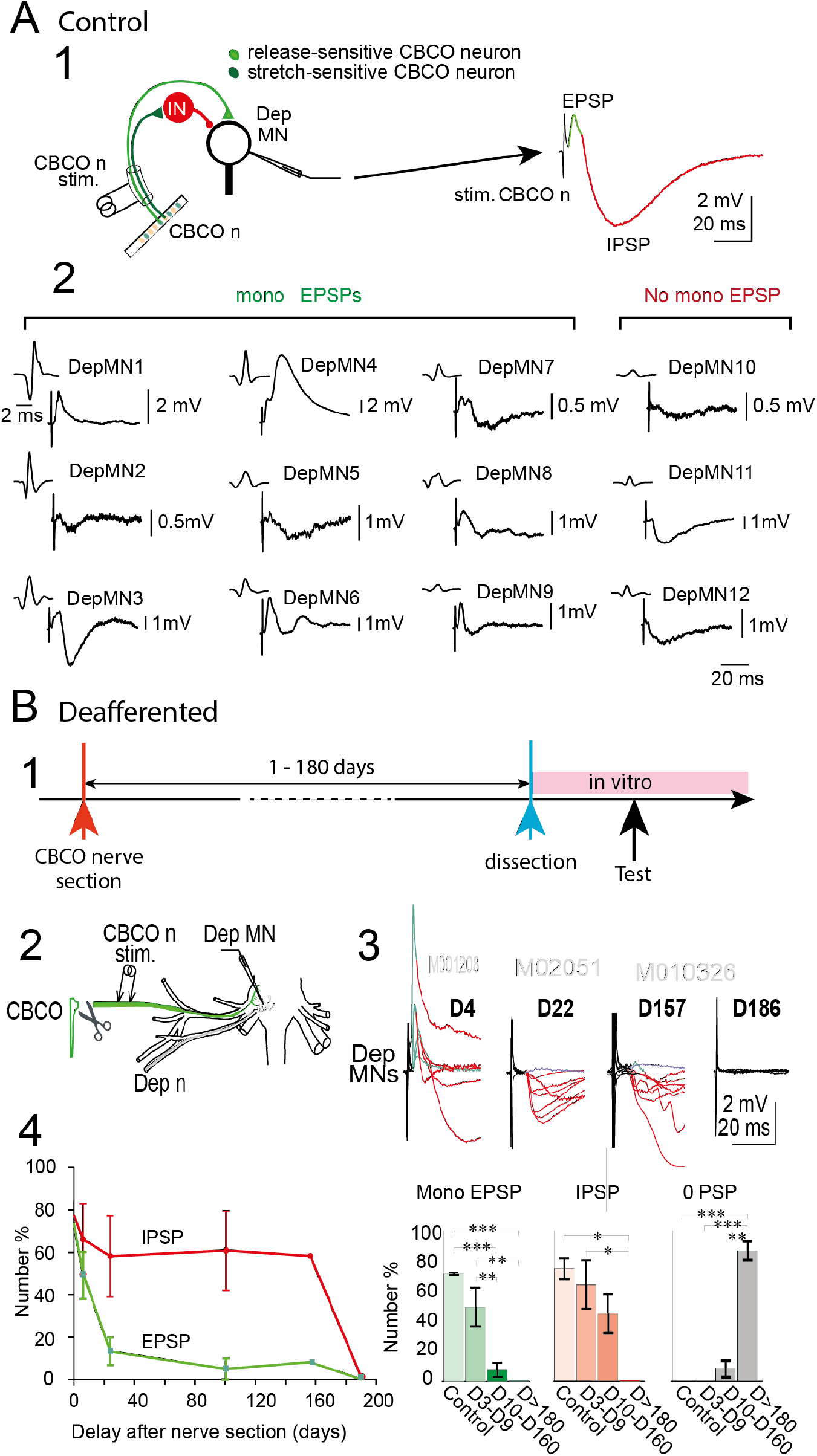
Evolution of synaptic transmission from CBCO to MNs after axotomy. **A:** In **control situation**, CBCO nerve stimulation (*CBCO n stim*.) evokes dual (*EPSP/IPSP*) responses in Dep MNs. This mixed response is due to simultaneous stimulation of stretch- and release-sensitive CBCO neurons (*A1*). While stretch-sensitive CBCO neurons produce monosynaptic EPSP on 8 of the Dep MNs (green pathway), release-sensitive CBCO neurons elicit disynaptic IPSPs in Dep MNs (orange CBCO neurons and red pathway). These results are illustrated by successive intracellular recordings of the 12 Dep MNs in a single experiment (*A2*). **B**: In **deafferented situation**, the MN responses to CBCO nerve (central part) electrical stimulation evolves differently for EPSPs and IPSPs along time. B1: Time-line of the experiment; B2: *In vitro* preparation of the sensorimotor system previously deafferented; B3: Intracellular recordings from all 12 Dep MNs made at D4, D22, D157 and D186 after CBCO cut *in vivo*; B4: percentage of MNs in which IPSPs (red trace) and EPSPs (green trace) were recorded. Whereas the percentage of MNs with IPSPs decreases from 80% to 60% within three weeks and then remains similar up to D157, the percentage of MNs with EPSP dramatically decreases from 80% to <20% in the same period of time. Note that at D186 all EPSPs and IPSPs have totally disappeared. At the 4 periods (control, D3-D9, D10-D160 and >D180 the observed decrease is significant for the % of MNs presenting EPSPs (*Mono EPSP*), IPSPs (*IPSP*) and the percentage of MNs without any response (*0 PSP*) (One Way ANOVA followed by Tukey’s Multiple comparison test); *:p<0.05; **: p< 0.01; ***: p< 0.001; ****: p< 0.0001.

### Restoration of lost synaptic transmission by electrical stimulation

Surprisingly, even after all synaptic transmission is lost 6 months after sensory nerve section, it was still possible to restore synaptic function by applying repetitive electrical stimulation (0.5 Hz) of the sensory nerve during one hour in the in vitro preparation (Fig. 5A1). After this period of stimulation, each electrical stimulus elicited an IPSP in the intracellularly recorded MN (Fig. 5A2, 5A3). In the following stimuli, polysynaptic EPSPs were progressively recruited (see the progressive weakening of IPSP peak, orange arrow in Fig. 5A2), and after 20 minutes, some monosynaptic EPSPs were also observed (see fast EPSP in Fig. 5A2, yellow arrow and Fig. 5A3). Synaptic transmission was recovered in most MNs (Fig. 5B1). However, the recovered monosynaptic responses were significantly delayed (6.07ms ± 0.33 instead of 4.01 ± 0.42 ms in control, p<0.001, Fig. 5B2), whereas the delay of inhibitory responses was unchanged (13.01ms ± 0.74 instead of 12.83 ± 1.50 ms in control, p=0.91, Fig. 5B3).

**Figure 5:**
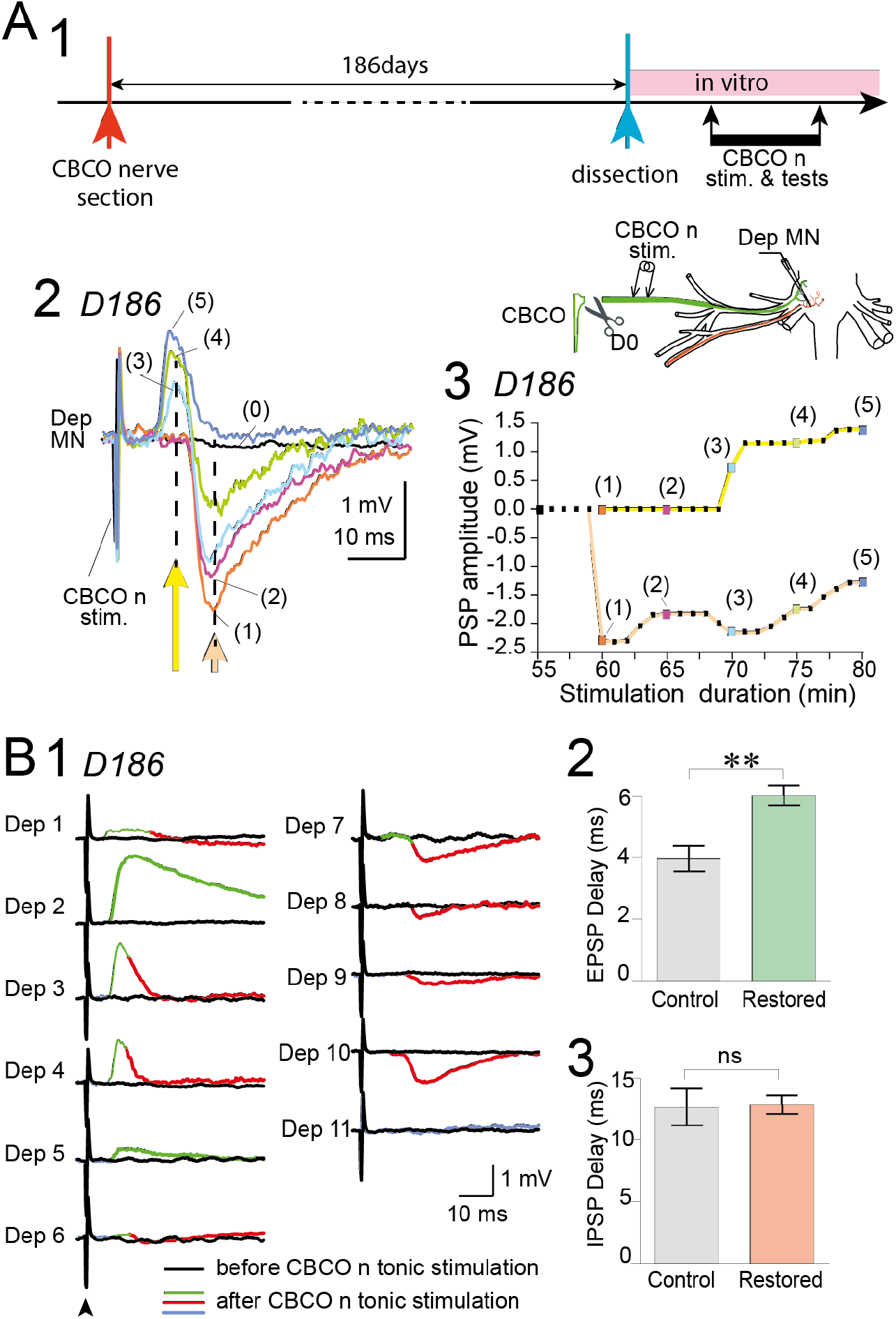
Restoration of monosynaptic EPSPs and disynaptic IPSPs in Dep MNs by electrical stimulation of the CBCO nerve *in vitro*. **A1** Timeline of the experiment. **A2** Results of an intracellular recording from a single Dep MN in a single experiment. The CBCO central part was tonically stimulated at 0.2 Hz. MN responses are superimposed over time (one record every 5 minutes). One can distinguish peak of IPSPs (*A2*, orange arrow) from peaks of EPSPs (*A2*, yellow arrow). **A3** The peak values of EPSPs and IPSPs are plotted along time. Note that IPSPs are restored before EPSPs. **B**: After the full restoration was observed (after 80 min stimulation), 10 other Dep MNs were successively recorded intracellularly. **B1**: Individual responses from the 11 Dep MNs (the first Dep MN recorded in A2, is labelled Dep 4 in B1). **B2-B3**: Comparison of synaptic delays measured for EPSPs (B2) and IPSPs (B3). Unpaired t-test: **: p<0.01; ns: p> 0.05.

### Spiking activity prevents loss of synaptic transmission

The loss of sensorimotor synaptic transmission described above was activity dependent. Indeed, when the sensory nerve was sectioned, chronic electrical stimulation (Fig. 6A1) applied *in vivo* to the proximal end (Fig. 6A1, B1-B3) was sufficient to avoid the loss of sensorimotor synaptic transmission (Fig. 6C, 6D) that was observed after 45 days (see Fig. 4B). In the *in vitro* preparations issued from these *in vivo* experiments, the electrical stimulation of the proximal end of the sensory nerve evoked an EPSP in 71.96%±3.22 of the recorded MNs (Fig. 6D1). This percentage was not significantly different from control preparations (74.24%±0.75), whereas in the absence of chronic stimulation, only 8.0%±4.9 of MNs displayed a monosynaptic EPSP. A similar tendency was also observed for IPSPs (Fig. 6D2) (82.27%±2.35 with chronic stimulation, 77.52%±7.11 in control, and 46.67%±13.32 without chronic stimulation) but the changes were not significant due to the lesser extend of IPSP loss.

**Figure 6:**
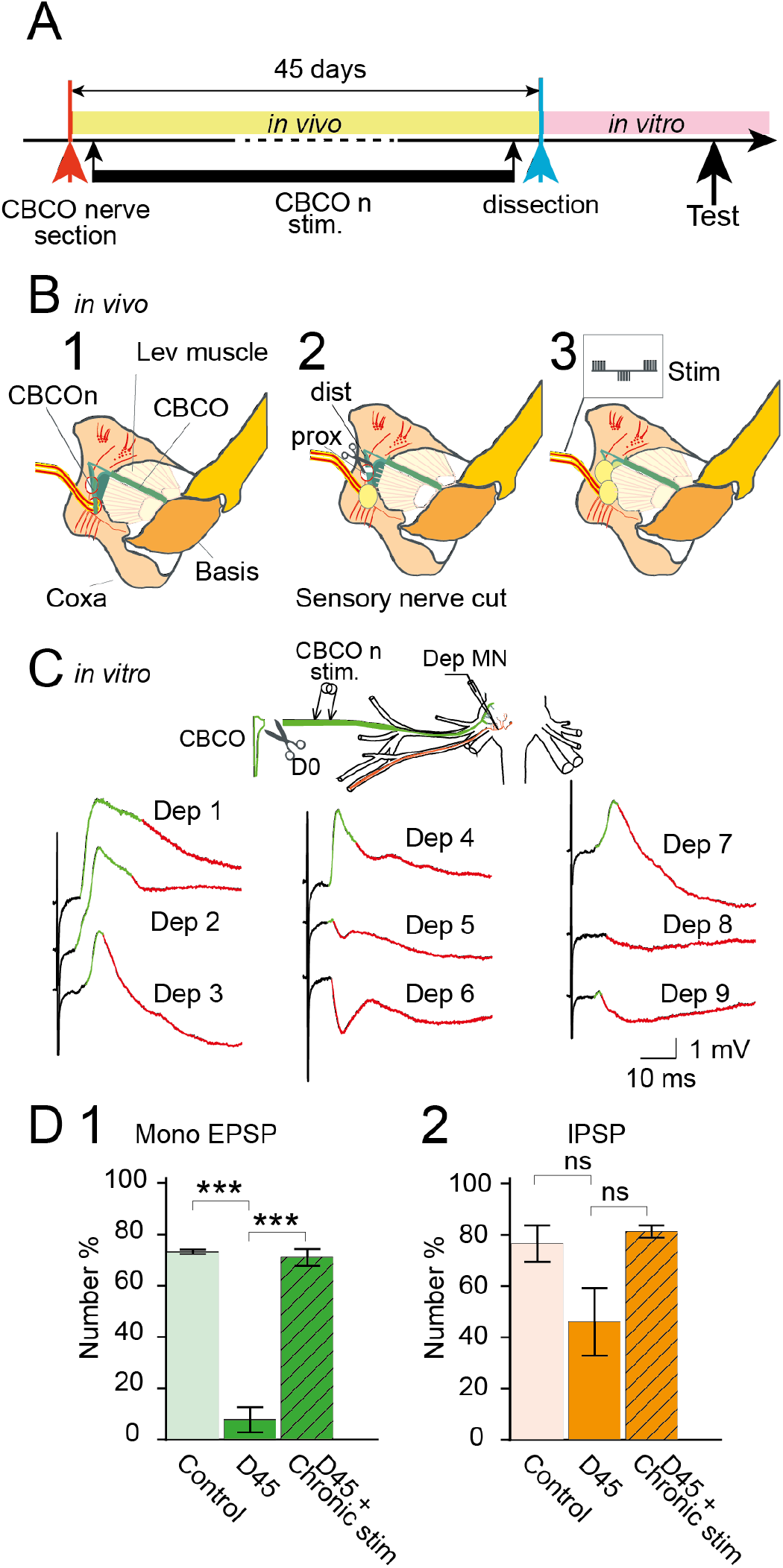
Prevention of PSP decline after CBCO cut. **A:** Timeline of the experiment. **B**: Experimental procedure for chronic stimulation of the central part of the cut CBCO. Two windows were made in the cuticle: the more central one was used to place two cuff electrodes on the CBCO nerve (*B1*); the more peripheral one was used to cut the CBCO nerve (*B2*). After the operation windows were covered with wax. Chronic electrical stimulation was applied to the cuff electrodes (*B3*). For more explanations see methods. **C**: Illustration of the results in one experiment. After 45 days, a dissection was made and the CBCO to Dep MNs pathways were studied in the *in vitro* preparation (see inset). Intracellular recordings from 9 Dep MNs were made in this experiment. Note that PSP were recorded in all of them. **D**: Statistical analysis of the % of Dep MNs presenting EPSPs (*D1*) and IPSPs (*D2*) in response to CBCO nerve stimulation *in vitro*. **D1**: Dep MNs presented an EPSP in 74.24% ± 0.76 in control situation (n=3), 8% ± 4.90 in CBCO cut + chronic stimulation situation (n=5) and 71.96% ± 3.22 in CBCO cut without chronic stimulation (n=3). ***: p< 0.001 (One Way ANOVA followed by Tukeys multiple comparison test). **D2**: Dep MNs presented an IPSP in 77.53% ± 7.11 in control situation (n=3), 46.67% ± 13.32 in CBCO cut + chronic stimulation situation (n=5) and 82.28% ± 2.35 in CBCO cut without chronic stimulation (n=3). ns: p> 0.05 (One Way ANOVA followed by Tukey’s multiple comparison test). Note that the observed decrease in the percentage of Dep MNs presenting an IPSP is not significant.

### Role of antidromic activity in preservation of CBCO output synapses

After peripheral section of the CBCO nerve, antidromic spikes were recorded in the proximal part of the nerve (connected to the CNS) (Fig 7). Given the anatomical disposition of output synapses from CBCO onto GABA interneurons (Watson *et al*., 2005), it is possible that such synapses disposed close to the main axon of a CBCO in the CNS (Fig. 2A-B), may be activated during PADs occurring slightly more distantly in a CBCO branch. This hypothesis would explain how, in the absence of sensory spikes, the reciprocal inhibitory pathway (Morgane Le Bon-Jego & Cattaert, 2002) could be activated by antidromic spikes (Fig. 2C, Fig.7), which would have preserved it, and would explain why it was easily restored even 9 months after the peripheral cut of the CBCO nerve. We tested this possibility by simulating of the involved circuits (Fig. 2C, Fig.8A-B) using NEURON 7.3 program (Hines & Carnevale, 1997), and by considering the spatial arrangement of output and input GABA synapses (Fig. 2D). This simulation confirmed that PADs that elicit antidromic spikes in the main CBCO axon are capable of recruiting output proximal synapses activating reciprocal inhibitory pathway evoking an IPSP in the MN (Fig. 8A-B), without evoking a monosynaptic EPSP in the MN.

**Figure 7:**
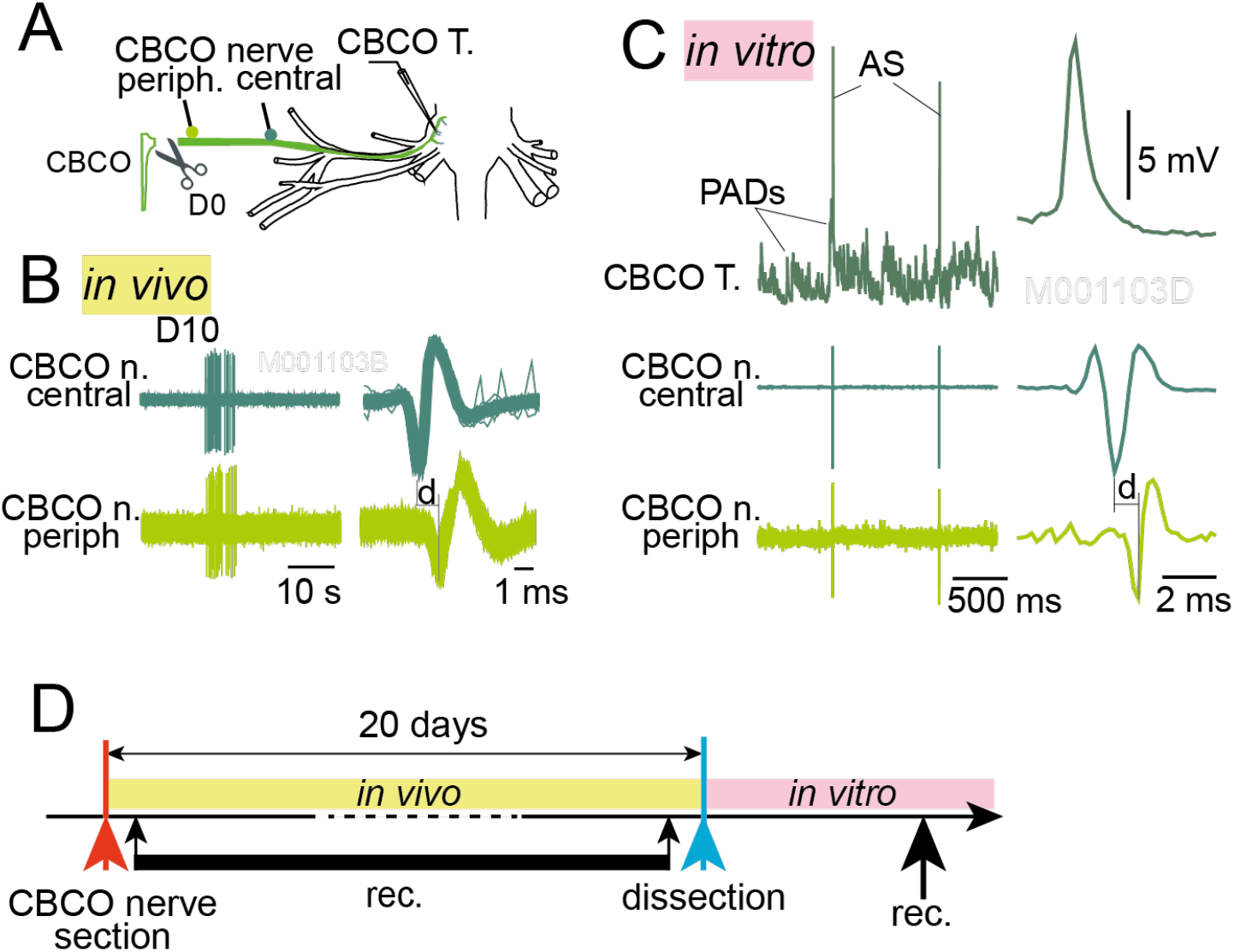
Antidromic discharges recorded in CBCO nerve *in vivo* and *in vitro* 20 days after CBCO cut. **A:** Experimental protocol: CBCO terminals (*CBCO T*.) were recorded intracellularly in the *in vitro* preparation and with en passant wire electrodes with two electrodes (peripheral and central). *In vivo*, two cuff electrodes were disposed on the CBCO nerve: one more peripheral and one more central. **B**: Result of *in vivo* extracellular recordings at peripheral and central sites of a CBCO nerve 20 days after CBCO cut. Note that the spike is recorded in the central site before the peripheral site, indicating that it is an antidromic activity. **C**: Result of *in vitro* recordings of another preparation dissected 20 days after CBCO section *in vivo*. Note that the spike is recorded first in the terminal branch of the CBCO neuron, then in the central en-passant electrode and finally in the peripheral en-passant electrode, indicating that this were antidromic spikes. **D**: Timeline of the above experiments.

**Figure 8:**
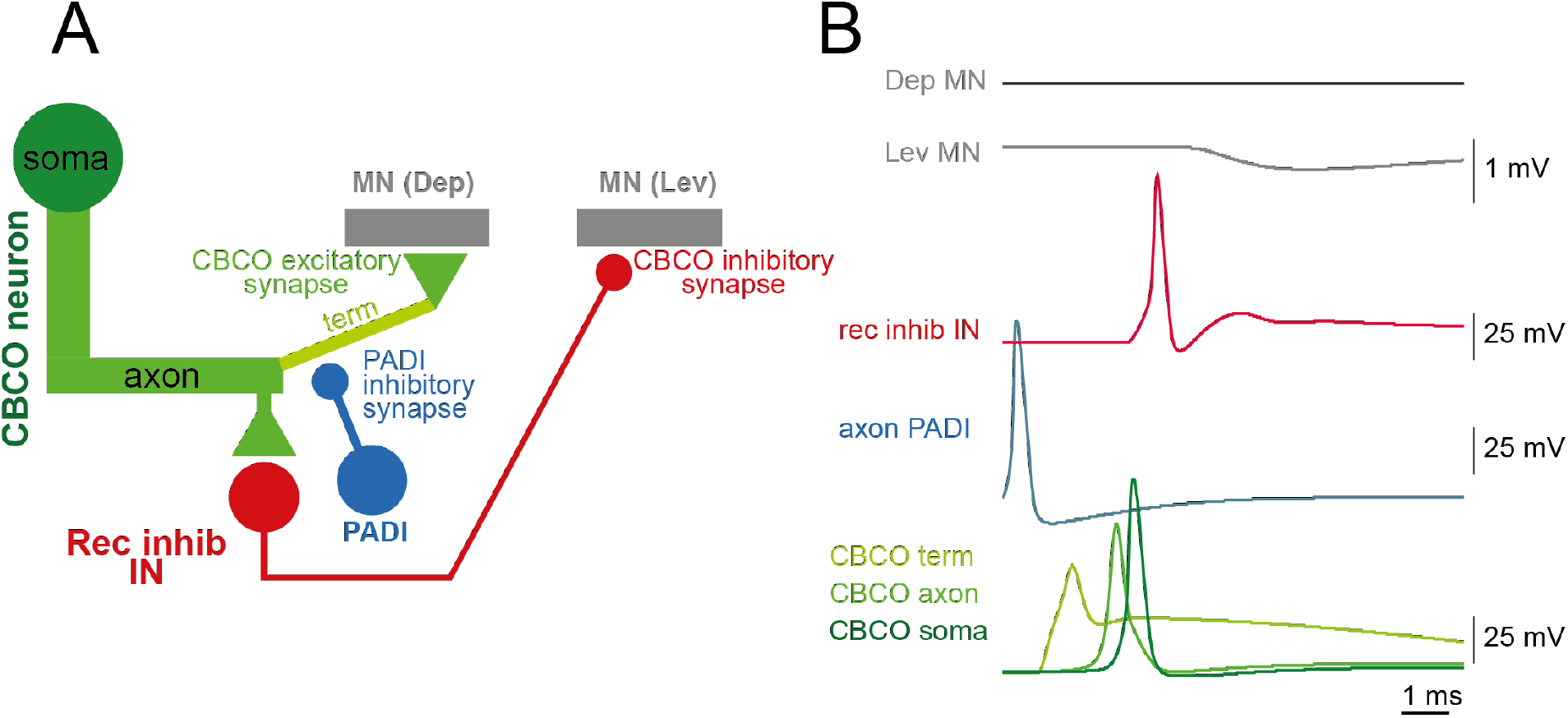
Simulation of the effect of PAD on CBCO-IN (reciprocal inhibition interneuron) synapse. **A:** Organization of the circuit modeled under Neuron script. The output synapse from primary afferent fibers (*CBCO axon*, in dark green) onto the *reciprocal inhibition GABA interneuron* (in red; connection observed in **A, B**) is slightly more peripheral than the *PADI* input synapses (in blue) onto the primary afferent (not shown, but see Watson et al., 2005). Two antagonistic motoneurons (MN) (in gray) receive an excitatory synapse from the primary afferent (*Dep MN*) and an inhibitory synapse from the reciprocal inhibitory IN (*Lev MN*), respectively. **B**: Result of simulation in which the *PADI* is activated and elicits a PAD in the primary afferent terminal (*CBCO term*) that triggers an antidromic spike recorded later in the primary afferent *axon* and *soma (CBCO axon* and *CBCO soma*, respectively). This depolarization is capable of eliciting an EPSP in the reciprocal inhibition interneuron (*rec inhib IN*) that triggers a spike, which is responsible for the IPSP observed in the *Dep MN* neurite. Note that the antidromic spike do not trigger any EPSP in the MN. This simulation demonstrates the possibility that PAD could activate the reciprocal inhibitory IN, and thereby could maintain the first synapse of the disynaptic inhibitory pathway from CBCO to MNs.

## Discussion

Our study outlines the absence of anatomical and functional degeneration in axons and their central synapses in neurons deprived of nucleus for long periods of time (> 6 months). Spike propagation and synaptic transmission persist within central nervous system in axotomized sensory neurons deprived of their cell bodies. This absence of degeneration of axon functionality is accompanied by a deep reorganization of glial cell population (proliferation and migration through endoneurium to get in close contact with the axons). Our study therefore confirms preceding findings about absence of anatomical and functional degeneration of sensory axons (Govind *et al*., 1992) and motor axons (Parnas *et al*., 1998) deprived of axon in crustacean. In both of these studies, proliferation and migration of glial cell nucleus in the axon tube were observed. Indeed, long survival of invertebrate axons deprived of their nucleus, and their capacity to conduct action potentials has been reported in invertebrates (Hoy *et al*., 1967, 1967; Nordlander & Singer, 1972; Wine, 1973; Bittner & Johnson, 1974; Ballinger & Bittner, 1980; Bittner, 1988; Atwood *et al*., 1989; Blundon *et al*., 1990, 1990; Bittner & Baxter, 1991; Parnas *et al*., 1991; Sheller & Bittner, 1992), and in vertebrates (Matsumoto & Scalia, 1981; Cancalon, 1982; Lubińska, 1982; Zottoli *et al*., 1987; Blundon *et al*., 1990).

Here, we were interested in the **activity-dependence of the synaptic function** of nucleus-deprived sensory axons. After 9-months nucleus deprivation, the stimulation of the peripheral part of the sensory nerve did not produce any response in post-synaptic motoneurons (Fig. 4). However, after one hour of stimulation post-synaptic responses were recorded in MNs, indicating that the machineries to conduct spikes and to activate functional synapses were still activable. Why was the first hour unsuccessful? It may be because spike conduction was blocked and the electrical stimulation restored this function, or it may be due to the synaptic machinery that was blocked. Indeed, previous observations in crayfish had shown that sensory nerve section did not immediately prevent the nerve activity (induced by electrical stimulation) to activate the target networks (Govind *et al*., 1992). However, the same stimulation did not evoke any response 3 weeks after the lesion but this was because of axon degeneration, and, therefore, synaptic function was not explored. On the other hand, decentralized motor axon controlling lobster deep abdominal extensors, continue to conduct spikes and evoke post-synaptic responses even one year after nerve cut (Parnas *et al*., 1991).

Due to the absence of sensory structure, no sensory activity could be recorded in the sensory nerve (Fig. 1A2). However, antidromic activity was recorded (Fig. 7). In freely moving animals, such antidromic activity is linked to the central control of sensory terminal activity via presynaptic inhibition produced by depolarizing GABA events in invertebrates (D. Cattaert *et al*., 1992; Daniel Cattaert & El Manira, 1999; Daniel Cattaert *et al*., 2001) and vertebrates (Dubuc *et al*., 1985; Vinay *et al*., 1999, 1999). It seems, however, that the antidromic activity fades along time, and it was rarely observed after 6-month nucleus deprivation. Here, we have shown that maintenance of output synapses on primary afferents is dependent on spiking activity because chronic stimulation of the central end of the cut CBCO nerve prevented the loss of synaptic transmission (Fig. 6). Note that the loss of PSP in postsynaptic target neurons was not due to spike conduction problems because after electrical stimulation of the sensory nerve, spikes could still be recorded in the main nerve trunk (data not shown). Although we do not explore the mechanisms of this activity-dependence of synaptic function, our results that such a control exist. Moreover, as a consequence, of differential effects of antidromic spiking activity on 1) excitatory synapses to MNs (monosynaptic resistance reflex circuit) and 2) disynaptic inhibitory connections to antagonist MNs (reciprocal inhibitory circuit) these two types of responses did not disappear at the same time (Fig. 4B3). Due to anatomical arrangement, synapses from primary afferents onto MNs are located à the ending of sensory axons, in an area where antidromic spikes are never recorded. Therefore, these synapses are not maintained by antidromic activity. By contrast, synapses from primary afferent onto inhibitory interneurons of the reciprocal inhibitory circuit are located in the region of the first branch of the main sensory axons within the central nervous system (Fig. 2A-C), where PADs can elicit spikes. This hypothesis was validated with a simulation pointing out this difference in spiking activity due to PADs, in the vicinity of these two types of synapses (Fig. 8AB). Antidromic activity was recorded in intact sensory nerves and central part of cut sensory nerves. Although this antidromic activity tend to decrease along months after lesion, the presence of spikes could explain the difference observed between PPSE and PPSI triggered by electrical stimulation of the central part of the CBCO nerve (Fig. 4B3). Nevertheless, since two synapses are involved in the disynaptic inhibitory circuit the question remains open as to know what occurs to the synapse between IN and MN. If PADs are strong enough to activate the IN, then the synapse from IN to MN could follow the same evolution. If it is not the case, then it is also possible that these INs can be activated by other neurons, as was shown for the 1a INs in vertebrates (Jankowska & Hammar, 2013).

We hypothesize that the differential effects of antidromic spikes (triggered by PADs) on monosynaptic EPSPs and disynaptic IPSPs could be the reason of the order of reappearance of PSP during repetitive CBCO nerve stimulation (Fig. 5). Since synapse to inhibitory interneurons had been regularly stimulated by antidromic spikes during the first months after nerve section, the degree of blocking machinery was less than the synapse to MNs that was not activated by antidromic spikes. This would imply that the **process of synapse blocking is somewhat gradual**. More work on activity-dependence of synapse maintenance in crustacean is needed to understand the involved mechanisms and explain its gradual nature.

